# Sequential ER stress and inflammatory responses are induced by SARS-CoV-2 ORF3 through ERphagy

**DOI:** 10.1101/2020.11.17.387902

**Authors:** Xiaolin Zhang, Ziwei Yang, Xubing Long, Qinqin Sun, Fan Wang, Pei-hui Wang, Xiaojuan Li, Ersheng Kuang

**Affiliations:** Institute of Human Virology, Zhongshan School of Medicine, Sun Yat-Sen University, Guangzhou, Guangdong, 510080, China; Key Laboratory of Tropical Disease Control (Sun Yat-Sen University), Ministry of Education, Guangzhou, Guangdong, 510080, China; Advanced Medical Research Institute, Shandong University, Jinan, Shandong 250012, China

**Keywords:** SARS-CoV-2, ORF3, ERphagy, ER stress, inflammatory response, ER apoptotic cell death

## Abstract

Severe acute respiratory syndrome coronavirus 2 (SARS-CoV-2) infections have resulted in a number of severe cases of COVID-19 and deaths worldwide. However, knowledge of SARS-CoV-2 infection, diseases and therapy remains limited, underlining the urgency of fundamental studies and drug development. Studies have shown that induction of autophagy and hijacking of autophagic machinery are essential for infection and replication of SARS-CoV-2; however, the mechanism of this manipulation and function of autophagy during SARS-CoV-2 infection remain unclear. In the present study, we identified ORF3 as an inducer of autophagy and revealed that ORF3 localizes to the ER and induces FAM134B-related ERphagy through the HMGB1-Beclin1 pathway. As a consequence, ORF3 induces ER stress and inflammatory responses through ERphagy and sensitizes cells to ER stress-induced cell death, suggesting that SARS-CoV-2 ORF3 hijacks ERphagy and then harms ER homeostasis to induce inflammatory responses through excessive ER stress. These findings reveal a sequential induction of ERphagy, ER stress and acute inflammatory responses during SARS-CoV-2 infection and provide therapeutic potential for ERphagy and ER stress-related drugs for COVID-19 treatment and prevention.

**Importance:** SARS-CoV-2 infection and replication require autophagosome-like double-membrane vacuoles. Inhibition of autophagy suppresses viral replication, indicating the essential role of autophagy in SARS-CoV-2 infection. However, how SARS-CoV-2 hijacks autophagy and the function of autophagy in the disease progression remain unknown. Here, we reveal that SARS-CoV-2 ORF3 induces ERphagy and consequently induces ER stress to trigger acute inflammatory responses and enhance sensitivity to ER stress-induced apoptosis. Our studies uncover ERphagy-induced inflammatory responses during SARS-CoV-2 infection and provide a promising therapeutic approach for treating SARS-CoV-2 infection and inflammatory responses in COVID-19 by manipulating autophagy and ER stress.

## Introduction

The pandemic COVID-19 outbreak has infected more than fifty million people and killed more than one million people worldwide, and the number of patients and deaths are still rapidly increasing. Effective treatments and therapy are still far from satisfactory; the development of specific drugs, therapy and vaccines is quite urgent. Severe acute respiratory syndrome coronavirus 2 (SARS-CoV-2) is the causative agent of COVID-19. As an emerging severe coronavirus, SARS-CoV-2 is highly homologous to two other severe coronaviruses, SARS-CoV-1 and MERS-CoV, which have caused two pandemic outbreaks in the past twenty years^1, 2^. SARS-CoV-2 infection in high-risk populations, such as older persons and persons with underlying diseases, often causes severe respiratory failure, sepsis and even multiple organ failure, accompanied by severe acute inflammation and excessive cytokine release known as a “cytokine storm” ^3–5^. To date, knowledge of SARS-CoV-2 infection and disease remains very limited.

Autophagy is a conserved physiological process in eukaryotic cells that separates and wraps intracellular waste, damaged organelles and invasive microorganisms into double-membrane vacuoles and eventually degrades them through lysosomes ^6, 7^. It is finely regulated through many core autophagy-related genes and several upstream signal cascades that are primarily initiated from amino acid and glucose deprivation, extracellular stimuli, intracellular damage and so on ^6^. Autophagy plays essential roles in cell activities and behaviors, mainly maintaining intracellular homeostasis, removing intracellular threats, repairing organelle damage and protecting cells from death ^8, 9^. However, excessive autophagy also hastens cell death ^10, 11^. While autophagy has exhibited great therapeutic potential in many diseases ^12^, safe and effective approaches in clinical applications still require prolonged development.

Autophagy usually eliminates invading viral infection and promotes antiviral responses; however, autophagy can also be hijacked by viruses to facilitate infection, replication and pathogenesis ^13, 14^. Many enveloped viruses often enter, replicate, mature and transport through autophagy-related vesicles ^15–17^. Thus, autophagy inhibition by genetic approaches or inhibitors effectively suppresses viral infection and diseases ^18, 19^. Therefore, autophagy has become a promising therapeutic target for viral infectious diseases.

Coronaviruses differentially regulate autophagy through distinct mechanisms. For example, several viral proteins of SARS-CoV-1 and MHV induce autophagy for viral infection and replication ^20–23^, whereas MERS-CoV Nsp6 inhibits autophagy ^22^. However, autophagy inhibition by different drugs and inhibitors effectively blocks their infection and replication, and coronaviruses likely require low levels of basic autophagy for viral entry and incomplete autophagy for virion assembly and maturation.

Previous studies have shown that the autophagy inhibitors CQ and HCQ significantly suppress SARS-CoV-2 infection and replication ^24, 25^, leading to hopes that COVID-19 might be defeated by modulating autophagy. However, ineffectiveness or no observed benefit of CQ and its derivatives has been confirmed in clinical trials of COVID-19 therapies ^26–28^, indicating that autophagy-related therapeutic strategies are still far from clinical application. Importantly, the infection and replication of SARS-CoV-2 requires activation of autophagy, and autophagy-like double-membrane vesicles are employed for viral RNA export and viral replication ^29^. Therefore, elucidating the fine regulation of autophagy by SARS-CoV-2 and its function during infection and replication is quite urgent for understanding SARS-CoV-2 infection and pathogenesis and for developing autophagy-related therapies and drugs for COVID-19 treatment. In the present study, we performed systemic screening and found that SARS-CoV-2 ORF3 induces ERphagy through the HMGB1-Beclin1 axis, consequently activating ER stress and inflammatory responses through ERphagy and sensitizing cells to ER stress-related cell death. Our findings reveal a novel induction of autophagy, ER stress and inflammatory responses by SARS-CoV-2 and provide a potential therapeutic target for COVID-19 treatment and drug development.

## Results

### A systemic screening identified that ORF3 induces autophagy

To investigate how SARS-CoV-2 regulates autophagy, systemic screening of a viral protein-expressing library was performed using the Gaussia luciferase-based autophagy reporter actin-LC3-DN, which detects the cleavage of pro-LC3, a key precursor of the autophagic protein LC3/ATG8 ^30^. After two rounds of screening, we identified several viral proteins that may decrease autophagy-dependent pro-LC3 cleavage; comparatively, ORF3 increases autophagy-dependent pro-LC3 cleavage in a dose-dependent manner (Figure S1). To confirm that ORF3 serves as an inducer of autophagy, an ORF3-expressing plasmid was transfected into cells without or followed by HBSS starvation. The shift of LC3-I to LC3-II was obviously enhanced, while the p62 level was decreased by ORF3 expression; both were augmented under HBSS starvation (Figure 1A). Furthermore, we detected the activation of one key kinase, Akt, which negatively regulates autophagy, and found that Akt phosphorylation was greatly decreased by ORF3 expression, similar to HBSS starvation (Figure 1B). To further visualize the autophagosome in the absence or presence of ORF3 expression, the ORF3-expressing plasmid was cotransfected with GFP-LC3- or ptfLC3-expressing plasmid. Increased GFP-LC3 puncta were observed in the presence of ORF3 expression compared with empty vector transfection (Figure 1C), and the formation of LC3-positive autophagosomes (yellow puncta) and the maturation of autolysosomes (red puncta) were similarly enhanced by ORF3 expression (Figure 1D). Similarly, by electron microscopy, increased double-membrane vacuoles of autophagy were observed in ORF3-expressing cells compared with few autophagic vacuoles in empty vector-transfected cells (Figure 1E). These results suggest that ORF3 overexpression significantly induced autophagy.

**Figure 1.**
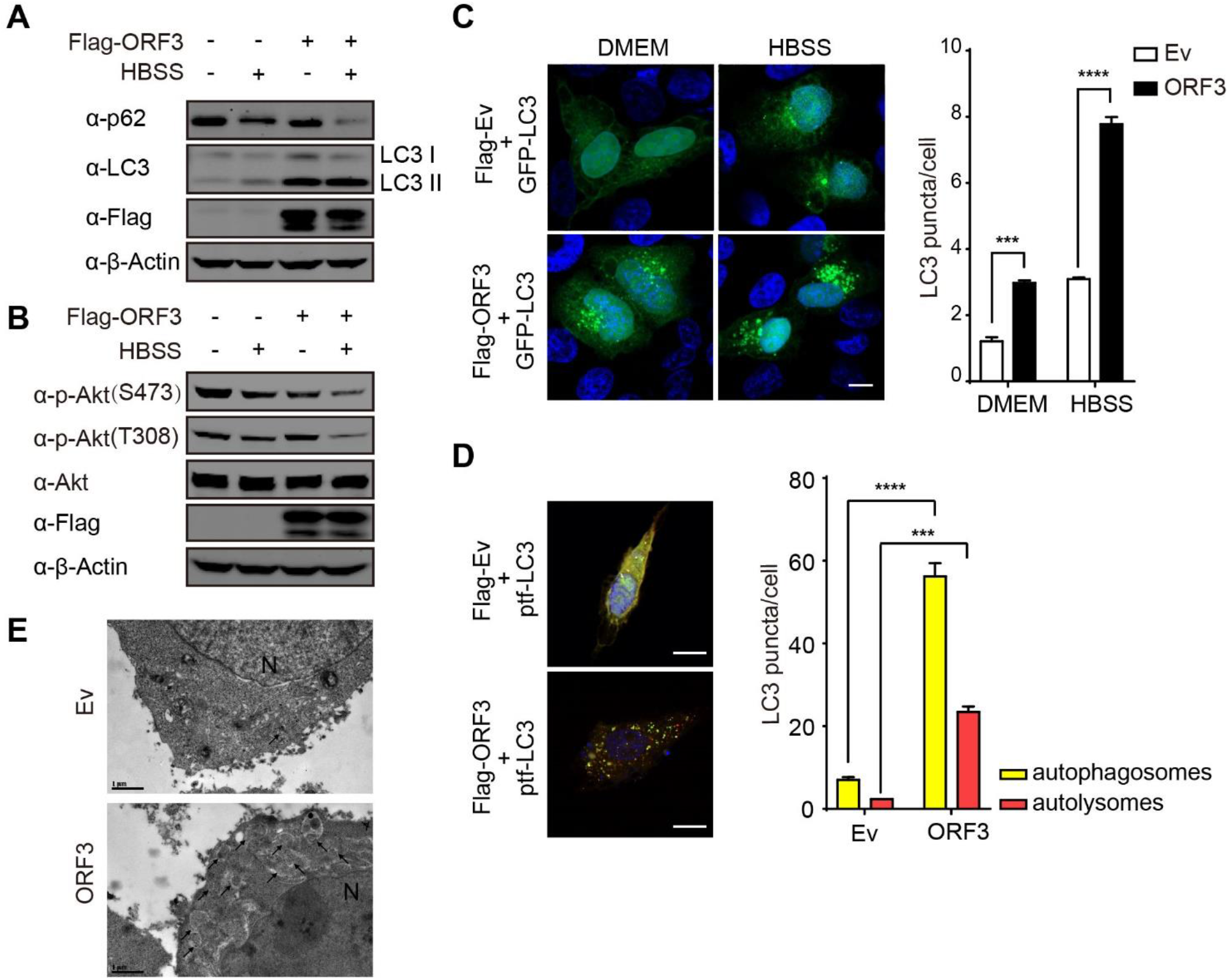
ORF3 induces autophagy. A-B. HEK293T cells were transfected with empty vector or Flag-ORF3-expressing plasmid for 48 h and then left untreated or treated with HBSS for 2 h. The cells were harvested, and the whole cell extracts were analyzed by Western blots as indicated to detect LC3-I/II shifts and p62 levels (A) or to detect Akt phosphorylation (B). C. HeLa cells were co-transfected with GFP-LC3 and empty vector or Flag-ORF3-expressing plasmid, left untreated or followed by HBSS treatment for 2 h, and then fixed and subjected to confocal microcopy analysis. Scale bar, 5 μm. The LC3 puncta per cell were calculated from 20 cells/each from three independent experiments. *, p<0.05, **, p<0.01, ***, p<0.001, ****, p<0.0001, Student’s t-test. D. HeLa cells were transfected with ptfLC3 and empty vector or Flag-ORF3-expressing plasmid. Twenty-four hours post transfection, cells were fixed and subjected to confocal microcopy analysis. Scale bar, 10μm. The LC3 puncta per cell were calculated from 20 cells/each from three independent experiments. The yellow puncta indicate autophagosomes, and the red puncta represent autolysosomes) *, p<0.05, **, p<0.01, ***, p<0.001, ****, p<0.0001, Student’s t-test. E. Autophagic vacuoles in control or ORF3-expressing HEK293T cells were observed by electron microscopy analysis, and representative images are shown. The arrows indicate autophagosomes. N, nuclei. Scale bar, 1 μm.

### ORF3 localizes to the ER and induces FAM134B-mediated ERphagy

To investigate the type of autophagy induced by ORF3, the subcellular location of ORF3 was determined by immunofluorescence. We found that OFR3 colocalized with ER membrane protein reticulon 3 (RTN3) around the nuclei (Figure 2A) and low colocalization levels with the autophagosome marker LC3 was observed, suggesting that ORF3 localizes to the ER but not in autophagic vacuoles. Alternatively, low colocalization levels of ORF3 with lysosomal red dye LysoTracker was observed (Figure 2B). Then, we presumed that ORF3 regulates ER-related autophagy, and the receptor of ERphagy FAM134B was determined in the absence or presence of ORF3 overexpression. The expression of FAM134B was not affected by ORF3 expression (Figure S2), while colocalization of ORF3 and FAM134B was observed (Figure 2C). Interestingly, FAM134B puncta were induced in the presence of ORF3 expression compared with the few FAM134B puncta observed in cells in the absence of ORF3 expression (Figure 2C). These results suggest that ORF3 promotes FAM134B-related ERphagy.

**Figure 2.**
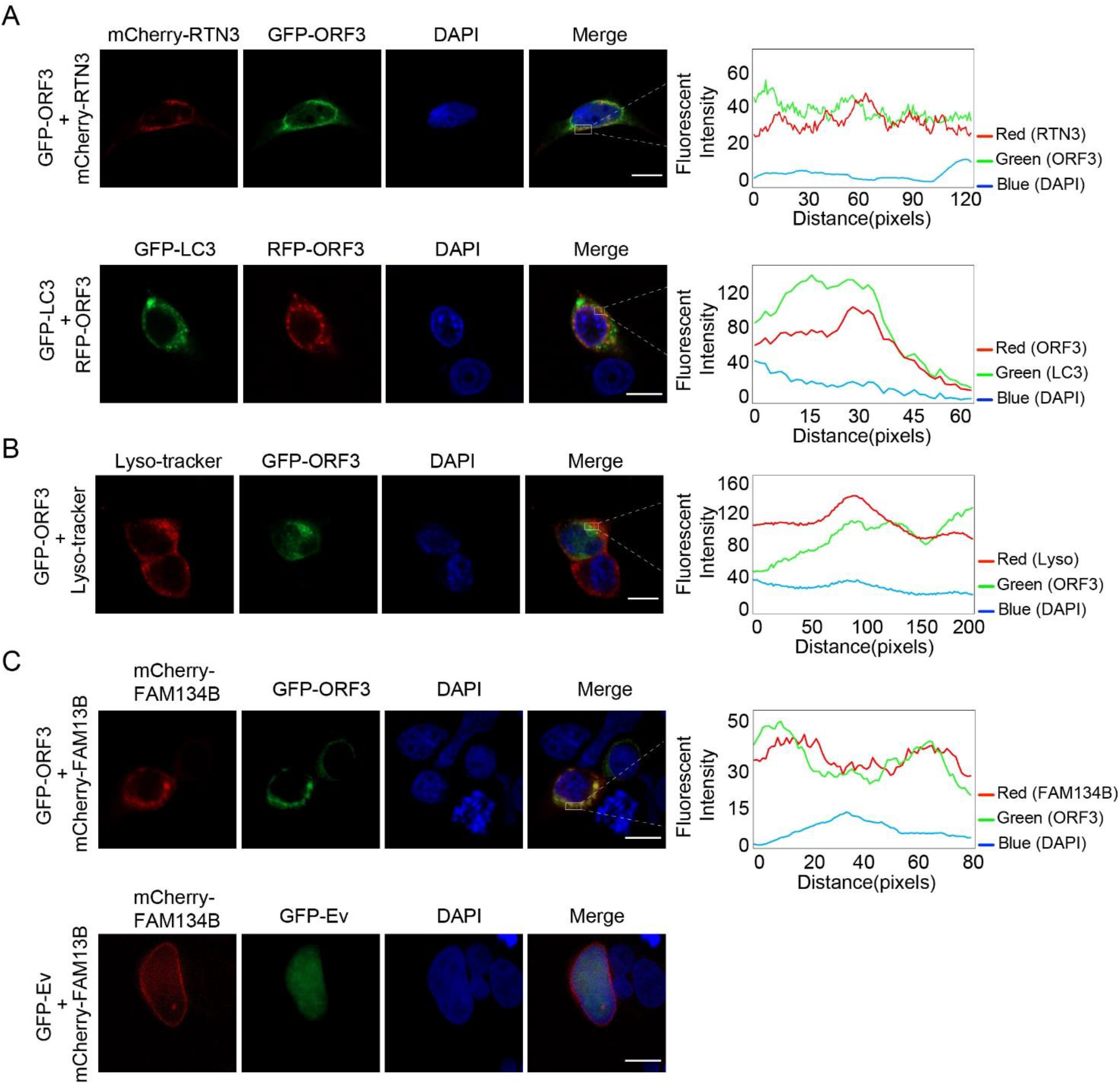
ORF3 localizes to the ER and induces FAM134B-mediated ERphagy. A. HeLa cells were cotransfected with GFP-ORF3 and mCherry-RTN3 or GFP-LC3 and mCherry-ORF3. Twenty-four hours post transfection, cells were fixed, and the images were visualized by confocal microcopy analysis. Scale bar, 5 μm. B. HeLa cells were transfected with GFP-ORF3 and then stained with the lysosomal red dye LysoTracker. The cells were fixed, and the images were visualized by confocal microcopy analysis. C. HEK293 cells were transfected with empty vector or GFP-ORF3- and mCherry-FAM134B-expressing plasmids. Twenty-four hours posttransfection, cells were fixed and subjected to confocal microcopy analysis. Scale bar, 5 μm. The coefficient of colocalization was determined by qualitative analysis of the fluorescence intensity of the selected area in Merge.

### ORF3 interacts with HMGB1 and induces Beclin-1-dependent autophagy

To investigate how ORF3 induces autophagy, we investigated public SARS-CoV-2 viral protein-binding protein network resources and found that several ORF3-binding proteins are involved in autophagy regulation ^31^. Five proteins were depleted by shRNA, and the effect on ORF3-induced autophagy was carefully detected. Depletion of high mobility group box 1 (HMGB1) and heme oxygenase 1 (HMOX1) greatly suppressed the ability of ORF3 to induce autophagy, whereas VPS11 and VPS39 depletion only weakly affected autophagy induction (Figure 3A and Figure S3A). As expected, the level of LC3-I-to-LC3-II shift induced by ORF3 was greatly decreased by HMGB1 shRNA in a dose-dependent manner (Figure 3B), confirming that HMGB1 is required for ORF3-induced autophagy. Furthermore, the interaction between ORF3 and HMGB1 was observed when both were transiently overexpressed in cells (Figure 3C). HMGB1 is associated with Beclin1 to enhance autophagy ^32, 33^, and the increased interaction of ORF3 with HMGB1 enhanced the association of Beclin1 with HMGB1 (Figure 3D), suggesting that ORF3 interacts with HMGB1 to enhance the HMGB1-Beclin1 association and subsequently induce autophagy. In addition, ORF3 interacts with HMOX1, and depletion of HMOX1 moderately attenuates the LC3-I to LC3-II shift in the presence of ORF3 expression (Figure S3B and S3C), indicating that HMOX1 may be involved in ORF3 induction in autophagy. Given that HMOX1 contributes to oxidative stress and HMGB1 acts as a key regulator of autophagy under oxidative stress, HMOX1 and HMGB1 may cooperate in the regulation of autophagy under oxidative stress in the presence of ORF3 expression.

**Figure 3.**
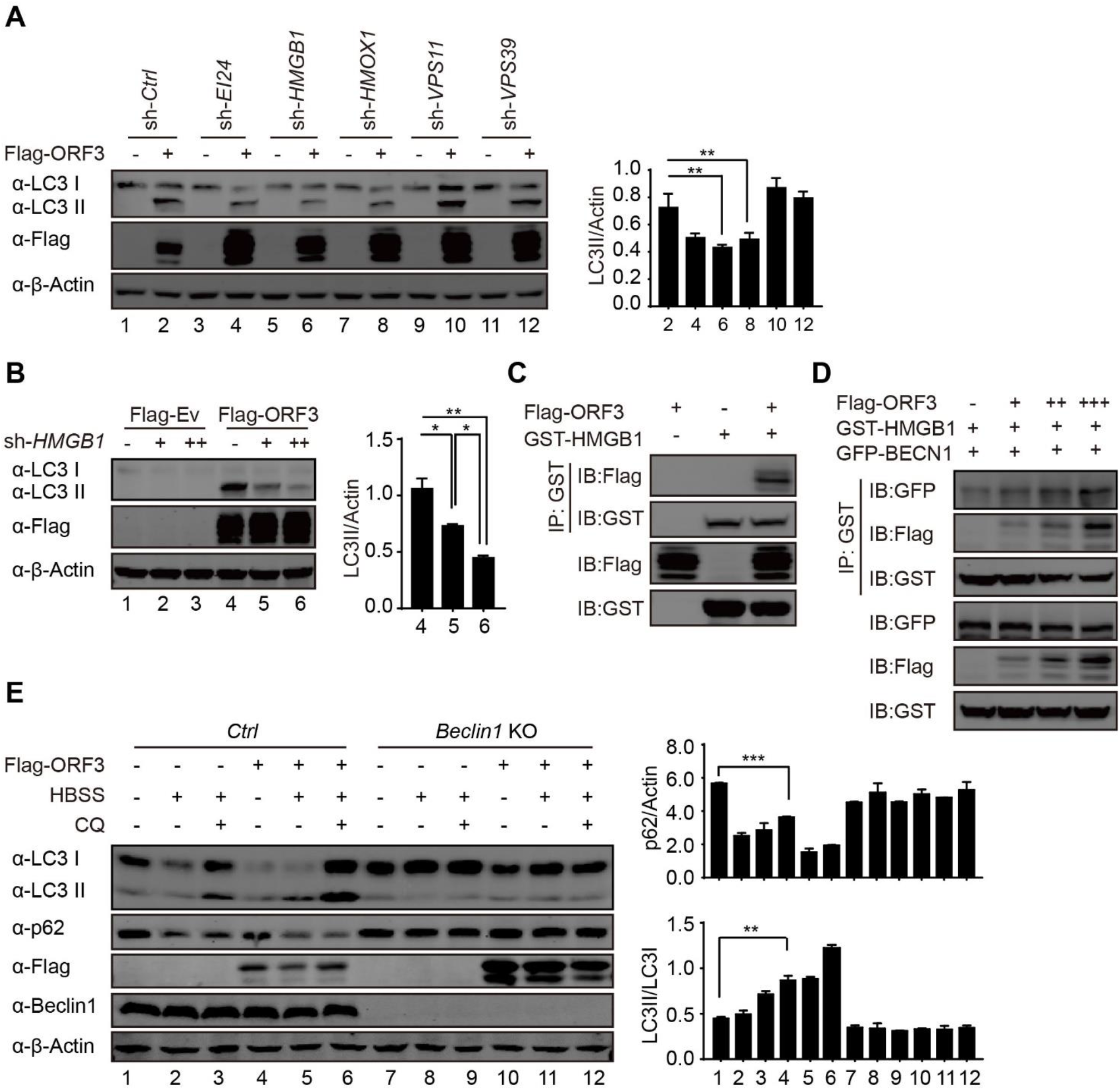
ORF3 interacts with HMGB1 and induces Beclin1-dependent autophagy. A. Flag-ORF3 or empty vector was cotransfected into HEK293T cells with scramble or shRNAs against EI24, HMGB1, HMOX1, VPS11 or VPS39. Thirty-six hours after transfection, cell pellets were collected, and the whole cell extracts were analyzed by Western blots as indicated to detect the LC3-I/II shift. The densities of LC3-II and actin bands were analyzed by a LI-COR Odyssey scanner and quantitated using ImageJ software. The relative ratio of LC3-II to actin was calculated from three independent experiments. The results are presented as the mean ± SD. *, p<0.05, **, p<0.01, ***, p<0.001, ****, p<0.0001, Student’s t-test. B. Scramble or shRNA against HMGB1 was transfected into HEK293T cells with Flag-ORF3 plasmid or empty vector. Thirty-six hours after transfection, cell pellets were collected, and the whole cell extracts were analyzed as indicated to detect the LC3-I/II shift. The relative ratio of LC3-II/actin was quantitated and analyzed as described above. C. GST-tagged HMGB1-expressing plasmid or empty vector was transfected into HEK293T cells with Flag-ORF3 or empty vector for 36 hours. Cells were collected, lysed and subjected to immunoprecipitation with GST-affinity beads, and then whole cell lysates and immunoprecipitated complexes were analyzed as indicated. D. Different amounts of Flag-ORF3-expressing plasmid or empty vector were transfected into HEK293T cells in the presence of GST-HMGB1- and GFP-Beclin1-expressing plasmids for 36 hours. Cells were collected and lysed, and immunoprecipitation was performed with GST-affinity beads. The whole cell lysates and immunoprecipitated complexes were analyzed as indicated. E. WT and Beclin1-KO HEK293T cells were transfected with Flag-ORF3-expressing plasmid and then left untreated or treated with HBSS for 2 h. Cells were harvested after an additional 50 μM chloroquine (CQ) treatment for 4 h, and then cell extracts were detected by Western blots with the indicated antibodies. The densities of LC3, p62 and actin were quantitated and analyzed using ImageJ software. The relative ratios of LC3-II to LC3-I and p62 to actin were calculated from three independent experiments. The results are presented as the mean ± SD. *, p<0.05, **, p<0.01, ***, p<0.001, ****, p<0.0001, Student’s t-test.

To further detect the role of Beclin1 in ORF3 regulation in autophagy, wild-type or Beclin1-deficient cells (Beclin1-KO) were transfected with ORF3-expressing plasmid, ORF3-induced increase of LC3-II/LC3-I shift and decrease of p62 level was completely attenuated in Beclin1-KO cells compared with that in wild-type cells (Figure 3E). These results suggest that ORF3 induces Beclin1-dependent autophagy through the HMGB1-Beclin1 pathway.

### ORF3 induces ER stress and inflammatory responses through ERphagy

To further investigate the function of ORF3-induced autophagy, the transcriptome of ORF3-expressing A549 cells was analyzed by RNA-sequencing analysis. The differentially expressed gene (DEG) clusters were enriched by GSEA analysis. ORF3-induced enriched DEGs shared high similarity to the DEGs in SARS-COV-2- and MERS-CoV-infected cells (Figure S4A). Interestingly, ER stress-related genes (Figure 4A, top) and innate immune and inflammatory genes (Figure 4A, middle and bottom) were enriched in the presence of ORF3 expression, indicating that ORF3 activates both ER stress and inflammatory responses. Alternatively, 97 DEGs were upregulated in both SARS-CoV-2-infected cells and ORF3-expressing cells (Figure S4B), and a cluster of genes involved in ER stress and immune and inflammatory responses were highly upregulated by both ORF3 expression and SARS-CoV-2 infection (Figure S4C), suggesting that ORF3 plays a critical role in the induction of ER stress and inflammatory responses during SARS-CoV-2 infection.

**Figure 4.**
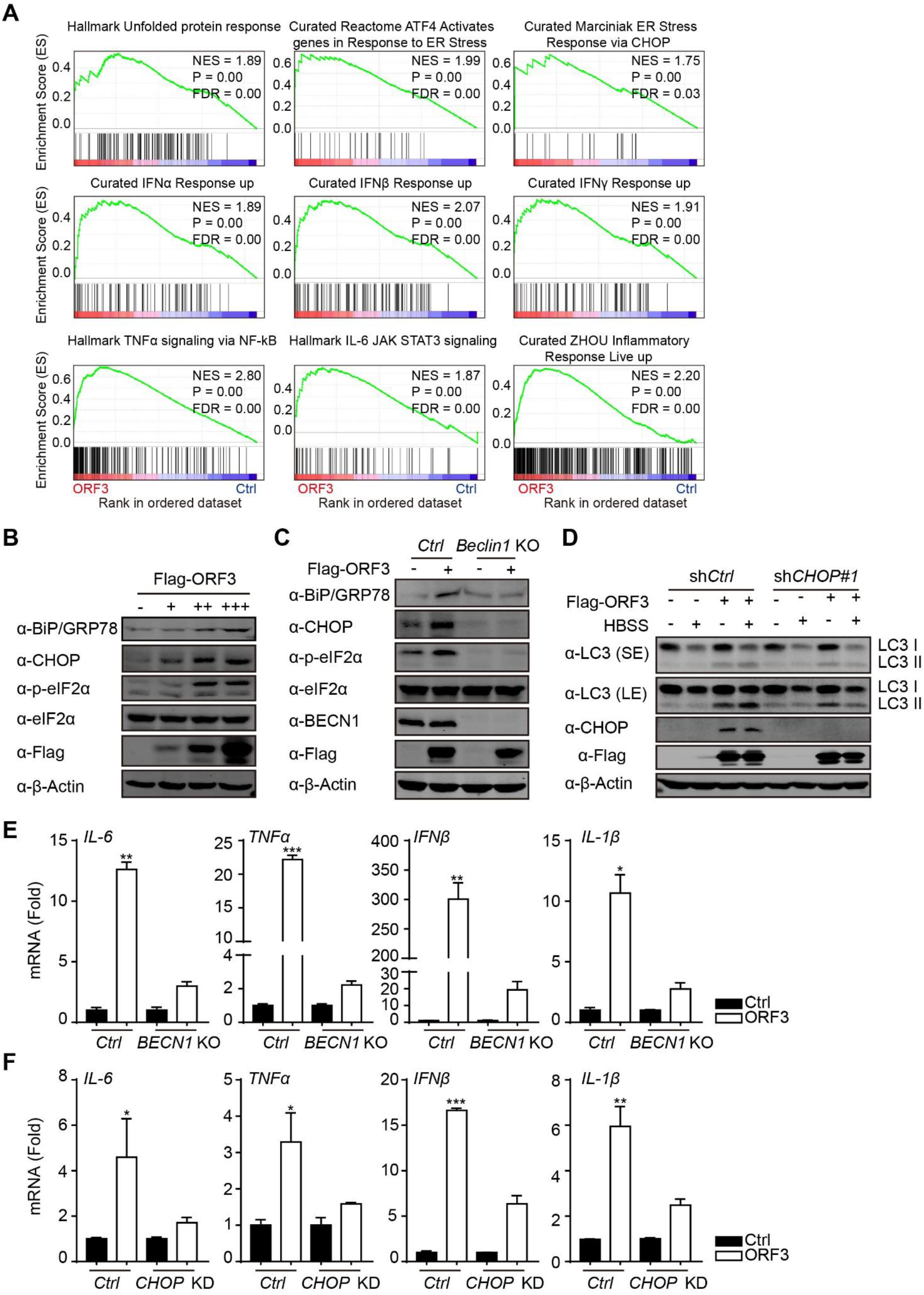
ORF3 activates ER stress and inflammatory responses through autophagy. A. GSEA analyses of RNA-sequencing data of ORF3-overexpressing A549 cells with “Hallmark gene sets” and “Curated gene sets”. NES, normalized enrichment score. FDR, false discovery rate. B. Flag-ORF3-expressing plasmid or empty vector was transfected into A549 cells. Thirty-six hours after transfection, cell pellets were lysed, and whole cell extracts were analyzed by Western blots with the indicated antibodies. C. Wild-type (WT) and Beclin1-KO HEK293T cells were transfected with Flag-ORF3-expressing plasmid for 36 hours, and then lysates were immunoblotted with the indicated antibodies. D. WT and CHOP-knockdown (CHOP-KD) A549 cells were transfected with Flag-ORF3-expressing plasmid or empty vector for 36 h, cell pellets were harvested after untreated or HBSS treatment for an additional 2 h, and then cell lysates were detected by Western blots with the indicated antibodies. E-F. WT and Beclin1-KO HEK293T cells (E) and WT and CHOP-KD A549 cells (F) were transfected with Flag-ORF3-expressing plasmid or empty vector as a control. Twenty-four hours post transfection, cells were treated with poly(I:C) (5 μg/ml) for 24 hours, and then total RNA was extracted and subjected to quantitative RT-PCR analysis to detect IL-6, TNF-α, IFN-β and IL-1β expression. The results are presented as the mean ± SD, biological replicates n = 3, *, p<0.05, **, p<0.01, ***, p<0.001, ****, p<0.0001, Student’s t-test.

To confirm that ER stress was induced by ORF3 expression, ER stress-related gene expression was detected in ORF3-expressing cells. The expression of BIP/Grp78 and CHOP gradually increased following increasing ORF3 expression, and the phosphorylation of eIF2α was similarly increased by ORF3 expression (Figure 4B). Notably, the induction of ER stress by ORF3 expression requires autophagy, and the increased BIP/Grp78 and CHOP expression and eIF2α phosphorylation were completely abolished in Beclin1-KO cells compared to wild-type cells (Figure 4C). However, depletion of the ER stress-related gene CHOP barely affected autophagy induction by ORF3 expression (Figure 4D), indicating that ORF3 induces autophagy independent of ER stress. These results suggest that ORF3 induces ER stress through autophagy induction, probably because ORF3-induced excessive ERphagy reprograms or disrupts ER homeostasis.

ER dysfunction often triggers innate immune and inflammatory responses; thus, the expression of key cytokines and the proinflammatory factors IFNβ, IL-1β, IL-6 and TNFα were induced by ORF3 expression. The induction of these factors was dependent on autophagy because Beclin1 deficiency in Beclin1-KO cells abolished the increased expression of these cytokines compared with that in wild-type cells (Figure 4E). Similarly, depletion of CHOP expression also greatly attenuated their increased expression in the presence of ORF3 expression (Figure 4F). These results suggest that ORF3 induces ER stress and subsequent inflammatory responses through ERphagy.

### ORF3-induced ERphagy and ER stress sensitize cells to ER apoptotic cell death

Considering that ORF3 induces ER stress, we presumed that ORF3 disrupts ER function and promotes cell death under ER dysfunction. ORF3-expressing A549 cells were treated with tunicamycin, an inducer of ER stress that causes an unfolded protein response and induces ER stress by inhibiting the N-linked glycosylation of GlcNAc phosphotransferase. Under tunicamycin treatment, the phosphorylation of the stress-related kinases p38-MAPK and SAPK/JNK was increased in the presence of ORF3 expression as well as c-Jun phosphorylation (Figure 5A, top). Importantly, the cleavage of ER-specific caspase-12 was increased by ORF3 expression under tunicamycin treatment (Figure 5A, bottom). These results suggest that ORF3 enhances ER apoptotic cell death under ER stress.

**Figure 5.**
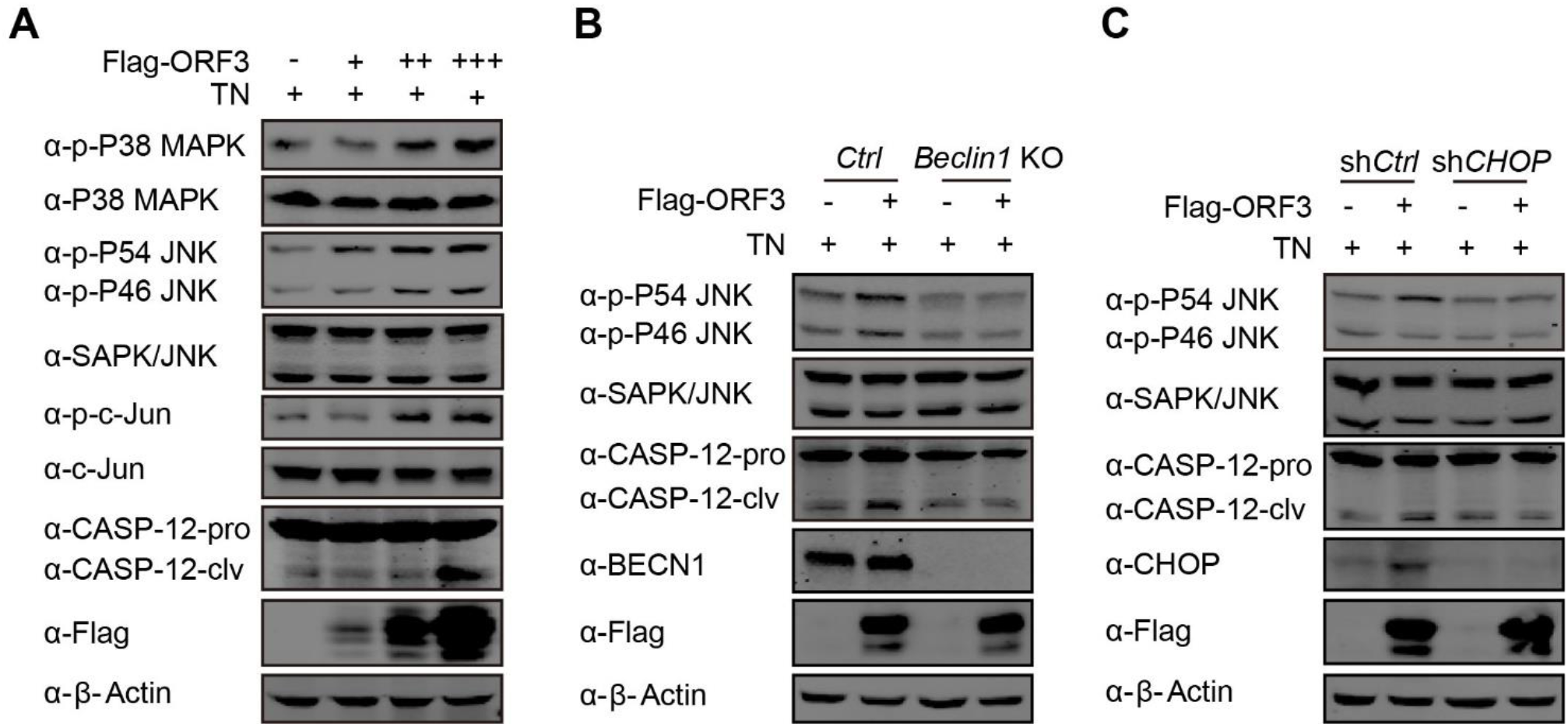
ORF3 enhances ER stress-induced apoptosis through autophagy. A. Flag-ORF3-expressing plasmid or empty vector was transfected into HEK293T cells. Twenty-four hours after transfection, cells were treated with 3 μg/ml tunicamycin (TN) for 15 h, cell pellets were collected, and whole cell extracts were analyzed by Western blots with the indicated antibodies. B. WT and Beclin1-KO HEK293T cells were transfected with Flag-ORF3-expressing plasmid, cells were harvested after 3 μg/ml TN treatment for 15 h, and then lysates were detected with the indicated antibodies. C. WT and CHOP-KD HEK293T cells were transfected with Flag-ORF3-expressing plasmid, cell extracts were prepared after TN (3 μg/ml) treatment for 15 h, and then lysates were detected by Western blots with the indicated antibodies.

To confirm that the increased cell death is responsible for ERphagy and ER stress, ER apoptotic cell death was further investigated under deficiency of autophagy or ER stress in Beclin1-KO or CHOP depleted cells, respectively. ORF3-induced phosphorylation of SAPK/JNK and cleavage of caspase-12 were greatly eliminated when ERphagy was blocked in Beclin1-KO cells (Figure 5B), indicating that ERphagy was required for ORF3-induced ER-related apoptosis. Similarly, the blockade of ER stress by CHOP depletion also eliminated ER-related apoptosis in ORF3-expressing cells (Figure 5C). Altogether, our results suggest that ORF3 overexpression triggers ER stress and ER abnormalities through ERphagy and consequently enhances the sensitivity of apoptosis to ER stress-related drugs.

## Discussion

Studies have confirmed that autophagy plays essential roles in the infection and replication of coronaviruses ^34, 35^; inhibition by genetic approaches and chemical inhibitors effectively suppresses their infection and might offer a promising therapeutic strategy for treating these infectious diseases. As an emerging severe coronavirus, the modulation and function of autophagy in SARS-CoV-2 infection and diseases remain unclear. By systemic screening, we revealed that SARS-CoV-2 ORF3 localizes to the ER compartment and induces FAM134B-mediated ERphagy through the HMGB1-Beclin1 pathway, subsequently contributing to ER stress and activating proinflammatory responses through ERphagy and ER functional reprogramming (Figure 6). As a consequence, sensitivity to ER stress-related drugs and ER apoptotic cell death are greatly enhanced in ORF3-expressing cells. This improves the therapeutic potential of autophagy- and ER stress-related drugs in the treatment of SARS-CoV-2 infection and COVID-19 disease for suppressing severe inflammatory responses and inducing cell death in SARS-CoV-2-infected cells.

**Figure 6.**
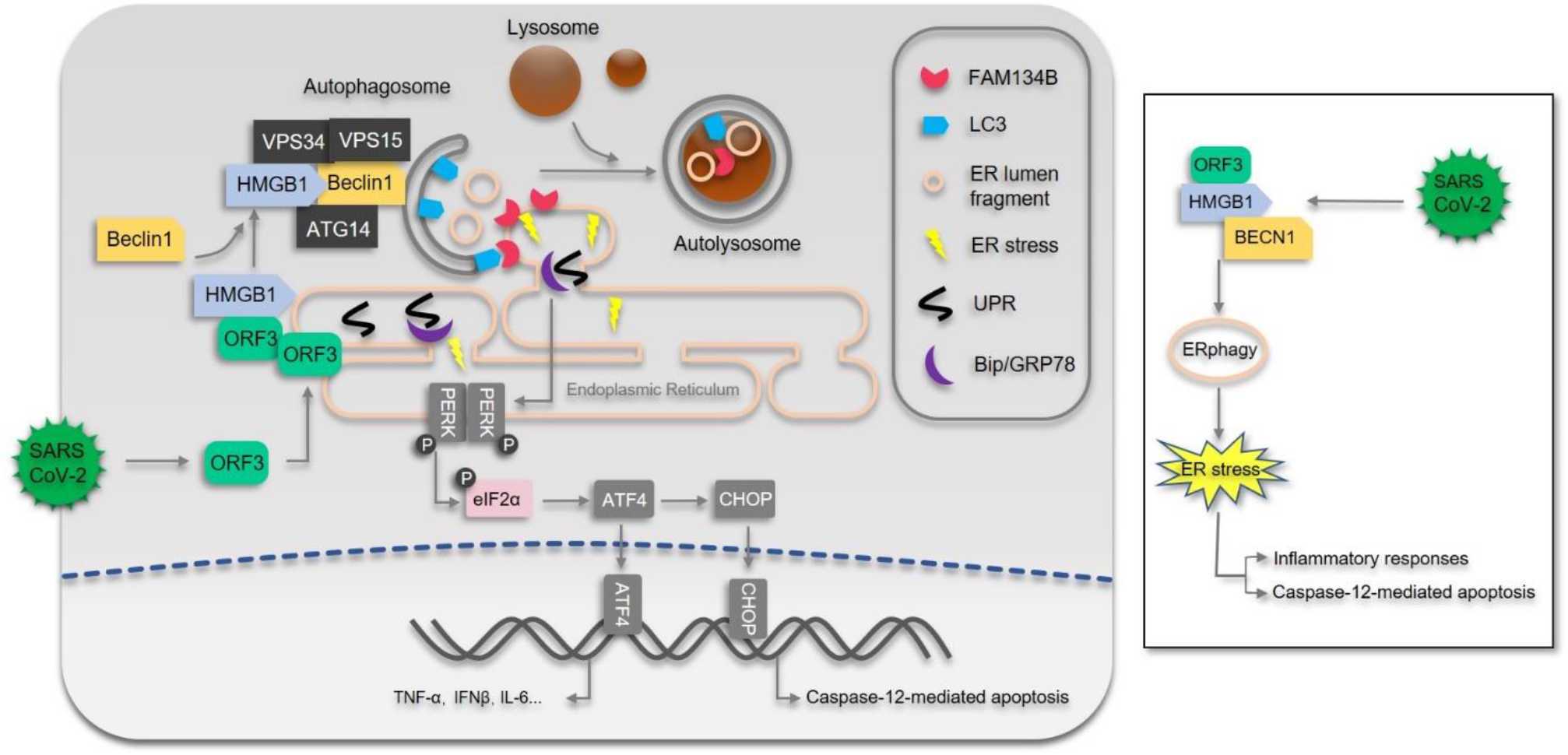
Diagram of ORF3-induced ERphagy, ER stress and inflammatory responses. During SARS-CoV-2 infection, ORF3 interacts with HMGB1 and enhances the association between HMGB1 and Beclin1 to initiate FAM134B-mediated ERphagy. Consequently, ER stress is induced by ORF3 through ERphagy to trigger proinflammatory responses and enhance the sensitivity of cells to caspase-12-mediated ER apoptotic cell death.

The autophagy inhibitor chloroquine and its derivative suppress SARS-CoV-2 infection and replication ^24, 25^, raising important hopes in therapeutic COVID-19 treatment and prevention. Although poor effectiveness and benefit have been concluded in clinical trials ^26, 27^, autophagy is confirmed to play essential roles in the SARS-CoV-2 life cycle. First, coronaviruses enter cells through endocytosis ^36^ and egress through lysosomes^37^, which require a fine control of autophagy and lysosomal degradation. Second, a recent study showed that SARS-CoV-2 infection and replication require autophagosome-like double-membrane vacuoles for RNA export and replication ^29^, indicating the essential requirement of autophagy and autophagic machinery for viral replication and assembly. Finally, the maturation and transport of virions of SARS-CoV-2 might also require autophagy or autophagic vacuoles like other RNA viruses ^16^. However, excessive autophagy is not beneficial to either cells or viruses because killing infected cells or clearing intracellular viral infection by strong autophagy induction is an effective antiviral defense. Therefore, autophagic processes are finely regulated to limit the appropriate strength and duration for facilitating cell survival and viral replication. Inhibitory viral proteins or other products of autophagy must exist during SARS-CoV-2 infection, as indicated in our preliminary screening; i.e., several viral proteins can suppress autophagy (Figure S1). In fact, several viral proteins of coronaviruses induce incomplete autophagy, and replication of coronaviruses utilizes the autophagic machinery while not requiring the complete genes and pathways of autophagy ^23, 38^. Thus, more viral regulation of autophagy and its function in SARS-CoV-2 infection require further investigation.

Our findings reveal that ORF3-induced ERphagy reprograms ER homeostasis and activates ER stress, probably to tolerate and adapt to the accumulation of viral proteins and recover ER function under virus-induced stresses. Many viral proteins of coronavirus rush to the ER ^39^ and form double-membrane vacuoles from the ER for viral replication ^16, 29^. We confirmed that ORF3 localizes to the ER and induces FAM134B-mediated ERphagy. Consequently, ORF3 induces ER stress through Beclin1-dependent autophagy, and Beclin1 deficiency abolishes the activation of ER stress in the presence of ORF3 expression, representing a novel induction of adaptive ER stress through ERphagy during SARS-CoV-2 infection. Therefore, ERphagy and ER stress are activated by SARS-CoV-2 through ORF3 to restrict the appropriate overload of viral proteins in the ER membrane and lumen, recycle overloaded viral proteins or repair ER damage.

As a consequence, ERphagy and ER stress induced by ORF3 reprogram ER function and activate inflammatory responses. Studies have shown that SARS-CoV-2 infection in diverse cells induces ER stress and activates proinflammatory responses ^40, 41^. Our studies confirmed that inflammatory responses are mainly induced by ORF3 through ERphagy-mediated ER stress, and blockade of ERphagy by Beclin1 KO or ER stress by CHOP depletion greatly eliminates the induction of key cytokines and inflammatory factors. This provides evidence that ERphagy and ER stress are the key upstream modulators of inflammatory responses during SARV-CoV-2 infection and may act as first-line precipitating factors of cytokine storms in COVID-19 patients, offering a novel target and approach to alleviating acute inflammatory responses during early SARS-CoV-2 infection through suppression of autophagy and ER stress.

Finally, considering that ER stress is activated and ER function is reprogrammed by ORF3 expression, we presumed that ORF3 may enhance the sensitivity of SARS-CoV-2-infected cells to ER stress-related drugs. Indeed, ORF3-expressing cells are sensitive to ER stress-drug tunicamycin treatment. The stress-related kinases p38-MAPK and SAPK/JNK were activated and ER apoptotic caspase-12 cleavage was enhanced by ORF3 in the presence of tunicamycin treatment. Correspondingly, they were abolished in Beclin1-deficient cells or CHOP-depleted cells separately, indicating that ORF3 enhances ER apoptotic cell death through excessive ERphagy and ER stress. These findings also support the therapeutic potential of ERphagy and ER stress in COVID-19 treatment, consistent with a hypothesis that ER stress is increased in COVID-19 patients and may benefit treatment ^42–44^.

In summary, our studies identified a novel viral inducer of autophagy during SARS-CoV-2 infection. We found that ORF3 localizes to the ER and triggers ERphagy through the HMGB1-Beclin1 pathway, consequently inducing ER stress and inflammatory responses. Our findings revealed a sequential induction of ERphagy, ER stress and inflammatory responses by SARS-CoV-2 ORF3 expression, facilitate insight into the hijacking of autophagy, reprogramming of ER function and activation of acute inflammatory responses during SARS-CoV-2 infection and pathogenesis, and offer a novel therapeutic target for COVID-19 treatment by suppressing autophagy and modulating ER stress.

## Materials and Methods

### Cells, antibodies and chemicals

HeLa, HEK293, HEK293T and Beclin-1 KO HEK293T (kindly provided by Dr Yang Du, Sun Yat-Sen University) cells were cultured in DMEM (Gibco) supplemented with 10% fetal bovine serum (FBS); human adenocarcinoma lung tissue-derived epithelial (A549) cells were cultured in RPMI 1640 (Gibco) medium with 10% FBS.

The following antibodies were used in this study: anti-LC3B (#3868), Beclin-1(#3495), Akt (#4691), p-Akt(Thr308) (#2965), p-Akt(Ser473) (#4060), GRP78/BiP (#3183), CHOP (#2895), p-eIF2α (#3398), eIF2α(#5324), p-p38 MAPK(Thr180/Tyr182) (#4511), p38 MAPK (#8690), p-SAPK/JNK(Thr183/Tyr185) (#4668), SAPK/JNK(#9252), p-c-Jun (Ser73) (#3270), and c-Jun (#9165) were purchased from Cell Signaling Technology (Beverly, MA, USA); anti-p62 (P0067) and Flag (F3165) were purchased from Sigma-Aldrich (St. Louis, MO, USA).; anti-FAM134B (DF12997) was purchased from Affinity Biosciences (Cincinnati, OH, USA); anti-Caspase 12 (55238-1-AP) was purchased from Proteintech Group (Rosemont, IL, USA). Goat anti-mouse IRDye680RD (C90710-09) and goat anti-rabbit IRDye800CW (C80925-05) were purchased from Li-COR (Lincoln, Nebraska, USA).

### Plasmids

The Flag-ORF3-expressing plasmid was kindly provided by the Pei-hui Wang laboratory (Shandong University). Plasmids for GFP-ORF3 and GFP-Beclin1 were constructed and subcloned into the pEGFP-C2 vector. Plasmids for GFP-LC3 and ptfLC3 were purchased from Addgene. Plasmids for GST-HMGB1 and mCherry-RTN3 were constructed and separately subcloned into the p-mCherry-C2 and pEBG vectors. To generate gene knockdown, shRNA target sequences were subcloned into the pLKO.1 vector between EcoRI and AgeI sites. The sequences of shRNAs are listed in supplementary Table S1.

### Immunoprecipitation and Western blotting analysis

In brief, one 10-cm dish of cells was transfected with 10-15 μg plasmid for 48 h. The cells were then collected and lysed in the presence of a protease inhibitor cocktail (Roche) and phosphatase inhibitors. For immunoprecipitation, the same amounts of cell lysates were precleaned and incubated with antibodies at 4°C overnight, and then protein complexes were precipitated with protein G-agarose, or the precleaned cell lysates were incubated with GST affinity beads at 4°C overnight. After washing five times, the immunoprecipitated complexes were separated by SDS-PAGE and subjected to immunoblotting analysis. For Western blotting, 40-60 μg protein of whole cell extracts per lane was separated by SDS-PAGE and transferred to membranes. The membranes were blocked in 5% dry milk, incubated with primary antibodies at 4°C overnight, and subsequently incubated with species-matched IRDye680- or IRDye800-labeled secondary antibodies for 2 h at room temperature. After 5 washes, images were visualized using a LI-COR Odyssey system.

### Real-time PCR

Total RNA was extracted using TRIzol reagent (Invitrogen) and reverse-transcribed using HiScript® III RT SuperMix (Vazyme). Real-time PCR was performed with a SYBR Green I Master Mix kit (Roche) and LightCycler® 480 system. The primer pairs are listed in supplementary Table S1.

### Immunofluorescence staining

The cells were fixed with 4% formaldehyde in phosphate-buffered saline (PBS) for 30 min, permeabilized with methanol for 10 min, and blocked with 2% bovine serum albumin in PBS for 30 min. Then, the cells were incubated with primary antibody overnight. After three washes with PBS containing 0.1% Triton X-100, the cells were incubated with Alexa 488- or 555-labeled anti-rabbit IgG antibodies (Invitrogen, Carlsbad, CA) for 1 h. The cells were counterstained with 4,6-diamidino-2-phenylindole (DAPI) (Sigma, St. Louis, MO) followed by three additional washes. The cells were mounted in antifade agent on glass slides and visualized with a confocal fluorescence microscope (Zeiss LSM800 microscopy with a 64× NA oil-immersion objective).

### Luciferase assays

A Gaussia luciferase-based reporter to detect pre-LC3 cleavage was used for the detection of autophagy activation^30^. Briefly, the Actin-LC3-DN reporter was transfected into HEK293T cells with either empty vector or Flag-ORF3-expressing plasmid in a 96-well plate. Twenty-four hours later, the cell supernatants were collected, and the secretion of Gaussia luciferase was measured using the reagent for the measurement of Renilla luciferase activity in a TriStar multimode reader. For each analysis, three independent experiments were performed in triplicate, and the mean ± SD was calculated.

### Electron microscopy

Flag-ORF3-expressing or control HEK293T cells were scraped, resuspended in DMEM and centrifuged in 1.5-ml centrifuge tubes. The medium was removed, and the cell pellets were fixed in 3% glutaraldehyde overnight at 4°C. The pellets were gently collected and placed in a specimen bottle containing 3% glutaraldehyde for fixation for another 24 h. After the fixed solution was discarded, the cells were rinsed with 0.1 M phosphate buffer for 1 h, washed 3-4 times, and fixed in 1% osmium tetroxide for 1 h. This was followed by another 3×10 min washes in 0.1 M phosphate buffer for 1 h. After dehydration through a series of ethanol washes from 30% to 2 washes in 100%, the cells were incubated in a mixture of Epon and propylene oxide (3:1 in v/v) for 90 min, embedded in a mold of 100% plastic, placed in a 70℃ oven and allowed to harden overnight (Epon 812). The cells were cut using a diamond knife to obtain ultrathin slices; the slices were placed on a hexadecimal copper grid (Electron Microscopy Science, H200-Cu). After drying for 2 h, transmission electron microscopy was performed on these grids using an FEI Tecnai G^2^ Spirit Twin electron microscope (13500×).

### RNA sequencing analysis and data availability

A549 cells were transfected with Flag-ORF3 or empty vector for 36 h, and total RNA was extracted with TRIzol following manufacturer procedures. Approximately 5 μg of total RNA was subjected to deep RNA sequencing. Libraries were constructed and sequenced on an Illumina platform, and 150 bp paired-end reads were generated. Data analysis was performed with Molecule Annotation System 3.0 (Annoroad Gene Technology Co., Beijing, China). The raw sequencing datasets generated in this study are available on the NCBI Gene Expression Omnibus (GEO) server under the accession number GSE158484. The RNA-sequencing data of SARS-CoV-2-infected cells were downloaded under the accession number GSE147507 from previously published analysis^45^.

### GSEA analysis

Gene set enrichment analysis was performed using GSEA software (https://www.gsea-msigdb.org/gsea/index.jsp) with number of permutations = 1000, permutation type = gene_set, enrichment statistic = weighted and metric for ranking genes = Signal2Noise. Enrichment outputs were considered significant and selected when NES ≥ 1, p < 0.05 and FDR < 0.25.

## Author contributions

X.Li. and E.K. initiated the concept. X.Z., Z.Y., X.Long., X.Li. and E.K. designed the experiments and analyzed the data. X.Z., Z.Y., X.Long., Q.S. and F.W. performed the experiments. P.W. provided the reagents. E.K. wrote the paper.

## Acknowledgments

We thank all the members of our laboratory for their critical assistance and helpful discussions. This work is supported by grants from the Natural Science Foundation of China (81671996 and 81871643) to E.K. and the Natural Science Foundation of China (81971928) to X.Li.

---

**Figure S1.**
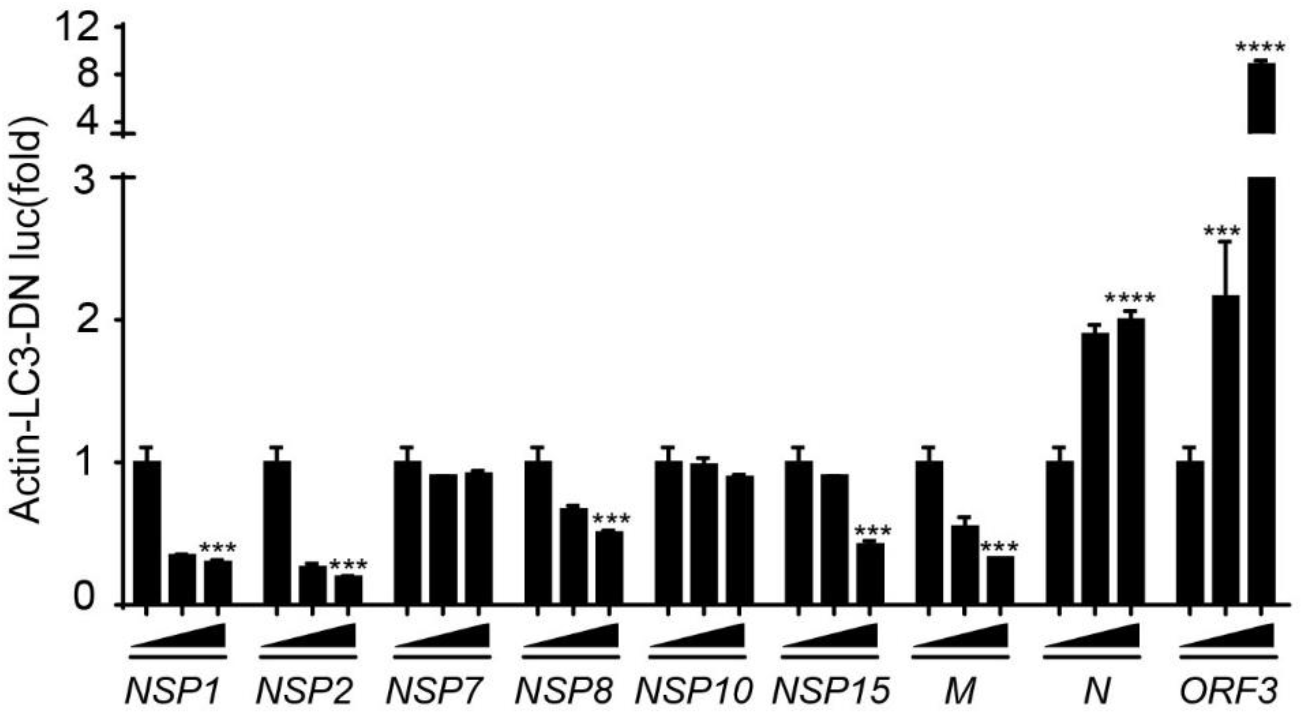
Screening results of viral proteins in autophagy regulation. HEK293T cells were transfected with empty vector or different amounts of NSP1-, NSP2-, NSP7-, NSP8-, NSP10-, NSP15-, M-, N- or ORF3-expressing plasmid plus Actin-LC3-DN-luc reporter. Twenty-four hours post transfection, the cells were removed, and then the supernatants were analyzed for actin-LC3-DN-luc activity. The results are shown as the mean ± SD, n = 3 biological replicates, *, p<0.05, **, p<0.01, ***, p<0.001, ****, p<0.0001, Student’s t-test.

**Figure S2.**
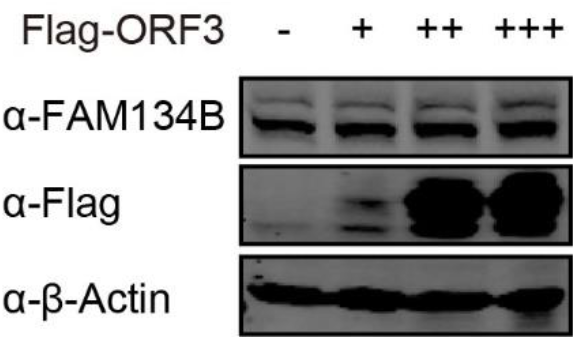
ORF3 does not affect FAM134B expression. Flag-ORF3 or empty vector was transfected into HEK293T cells. Thirty-six hours after transfection, cell pellets were collected, and the whole cell extracts were analyzed by Western blots to detect the level of FAM134B expression.

**Figure S3.**
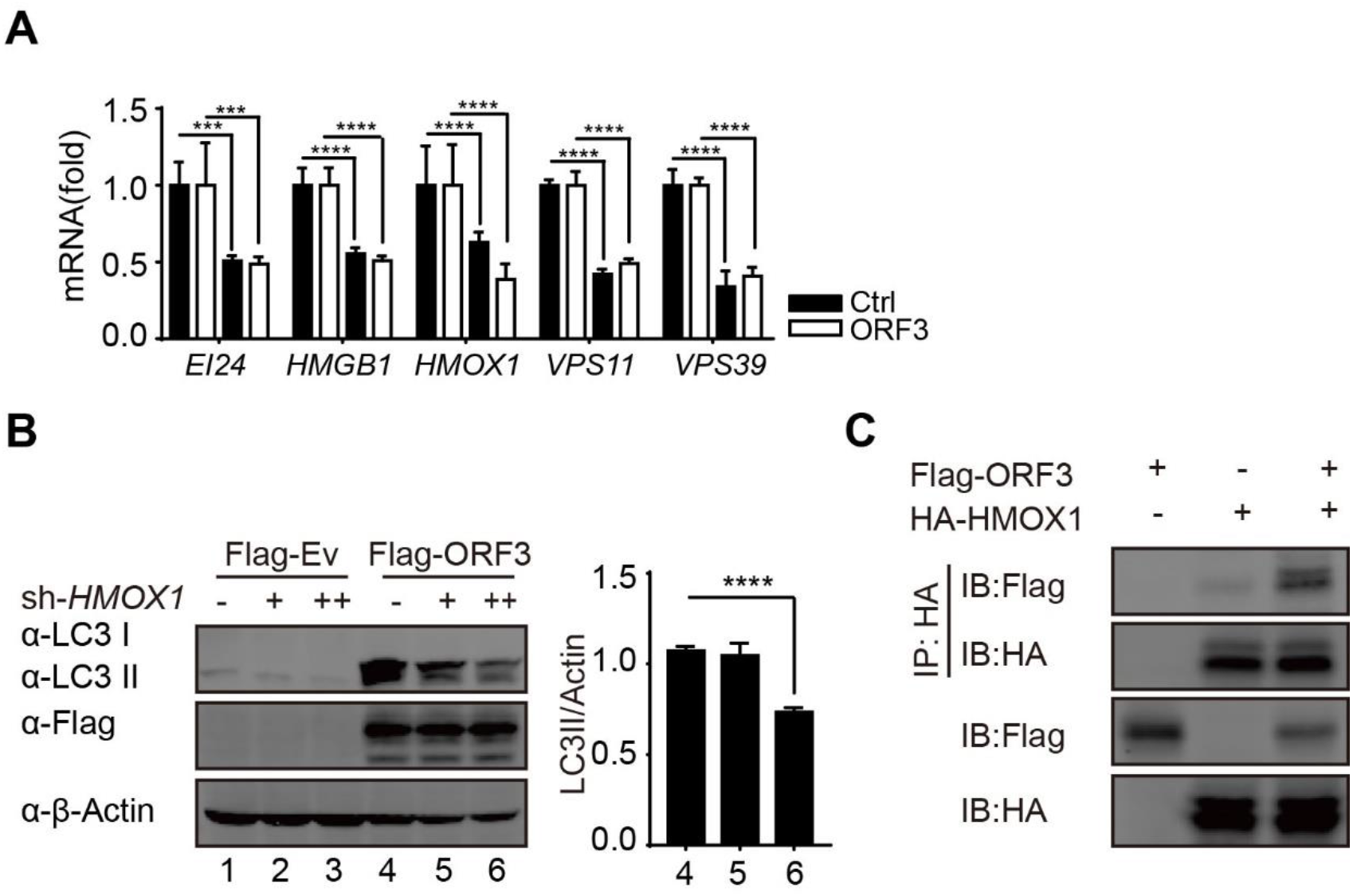
HMOX1 interacts with ORF3 and regulates ORF3-induced autophagy. A. Flag-ORF3 or empty vector was transfected into HEK293T cells with scramble or shRNAs against EI24, HMGB1, HMOX1, VPS11 or VPS39. Thirty-six hours after transfection, total RNA was extracted and subjected to quantitative RT-PCR analysis. The results are presented as the mean ± SD, n = 3 biological replicates, *, p<0.05, **, p<0.01, ***, p<0.001, ****, p<0.0001, Student’s t-test. B. Different amounts of shRNA against HMOX1 were transfected into HEK293T cells in the presence of Flag-ORF3-expressing plasmids or empty vector. Thirty-six hours after transfection, cell pellets were collected, and the whole cell extracts were analyzed by Western blots to detect the LC3-I/II shift. The densities of LC3-II and actin were visualized by a LI-COR Odyssey scanner and analyzed using ImageJ software. The relative ratio of LC3-II to actin was calculated for three independent experiments. The results are shown as the mean ± SD, n = 3 biological replicates, *, p<0.05, **, p<0.01, ***, p<0.001, ****, p<0.0001, Student’s t-test. C. HA-HMOX1 or empty vector was transfected into HEK293T cells with Flag-ORF3 plasmid or empty vector for 36 hours. Cells were collected, and immunoprecipitation with anti-HA-affinity beads was performed. Then, whole cell lysates and immunoprecipitated complexes were analyzed as indicated.

**Figure S4.**
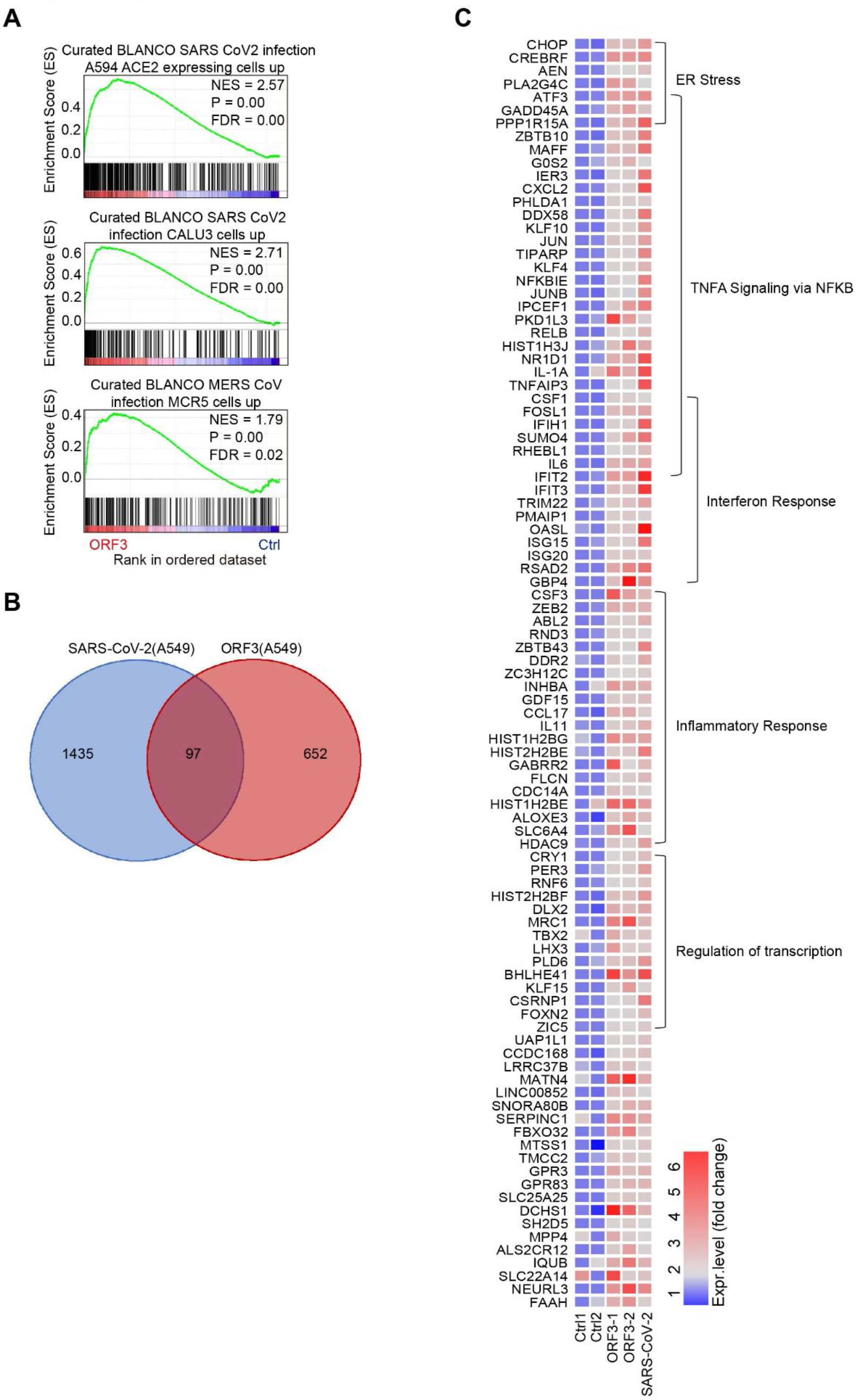
Differentially expressed genes in ORF3-expressing cells and SARS-CoV-2-infected cells. A. GSEA analyses of RNA-sequencing data of ORF3-expressing A549 cells with curated gene sets. NES, normalized enrichment score. FDR, false discovery rate. B. Venn diagram depicting upregulated genes shared between ORF3-transfected and SARS-CoV-2-infected A549 cells. C. Heatmap depicting the expression levels of the shared upregulated genes between the two groups, fold change >1.5, p <0.05.

**Supplementary Table 1.**
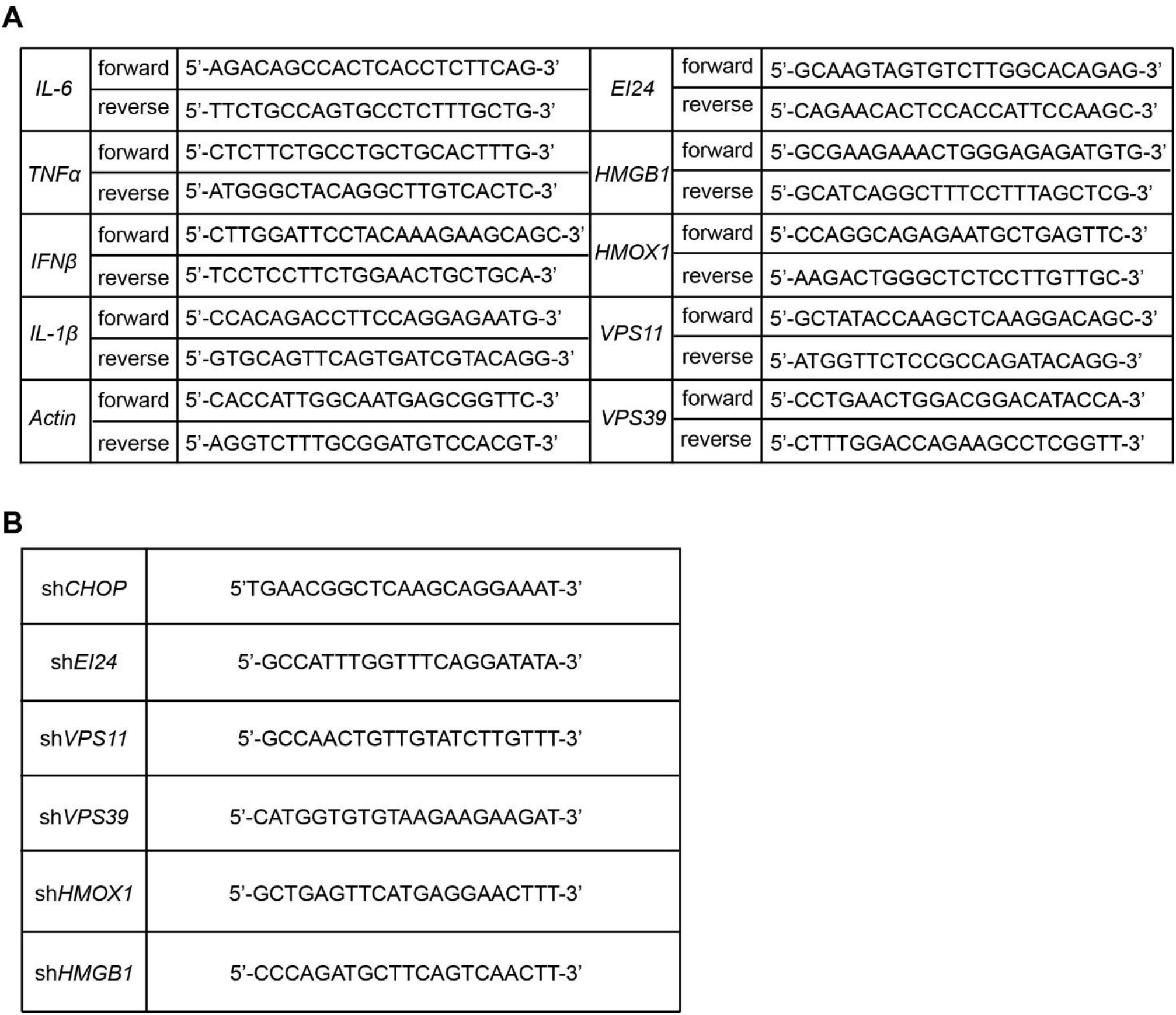
A. The sequences of primer pairs used in the RT-PCR array are listed. B. The target sequences of shRNAs used in this study are listed.

